## Supplementary File for "MR-SPLIT: a novel method to address selection and weak instrument bias in one-sample Mendelian randomization studies"

This supplementary file contains additional simulation and real data analysis results. The proof of Theorem 1 is given at the end of this file.

### 1 Results of selecting major IVs under different partial $F$ values

Table S1 shows the results of distinguishing major and weak IVs with different partial $F$ thresholds. We also showed the results with the criterion of $F > 100$. As demonstrated, this is exceedingly conservative and is generally best avoided. Furthermore, we recognize that a heritability ($h^2 = 0.5$) is considerably high for many exposure traits in practical scenarios, representing situations that are relatively uncommon in reality.

Table S1: Mean numbers of being identified as major IV using different criteria in 1000 simulations. For the noise category, it was aggregated over the 295 null IVs.

| $h^2$ | $N$ | Criteria | $SNP_1$ | $SNP_2$ | $SNP_3$ | $SNP_4$ | $SNP_5$ | Noises ($\times$295) |
|---|---|---|---|---|---|---|---|---|
| 0.15 | 500 | F>10 | 0.55 | 0.5 | 0.15 | 0 | 0.1 | 1.25 |
| | | F>30 | 0.05 | 0.05 | 0 | 0 | 0 | 0 |
| | | F>50 | 0 | 0 | 0 | 0 | 0 | 0 |
| | | F>100 | 0 | 0 | 0 | 0 | 0 | 0 |
| | 1000 | F>10 | 0.95 | 0.95 | 0.35 | 0.1 | 0 | 0.65 |
| | | F>30 | 0.35 | 0.35 | 0 | 0 | 0 | 0 |
| | | F>50 | 0.15 | 0 | 0 | 0 | 0 | 0 |
| | | F>100 | 0 | 0 | 0 | 0 | 0 | 0 |
| | 2000 | F>10 | 1 | 1 | 0.75 | 0.35 | 0.55 | 0.6 |
| | | F>30 | 1 | 0.95 | 0.1 | 0 | 0 | 0 |
| | | F>50 | 0.75 | 0.75 | 0 | 0 | 0 | 0 |
| | | F>100 | 0 | 0 | 0 | 0 | 0 | 0 |
| 0.3 | 500 | F>10 | 0.95 | 0.8 | 0.25 | 0.25 | 0.1 | 1.15 |
| | | F>30 | 0.5 | 0.55 | 0 | 0 | 0 | 0 |
| | | F>50 | 0.25 | 0.25 | 0 | 0 | 0 | 0 |
| | | F>100 | 0 | 0 | 0 | 0 | 0 | 0 |
| | 1000 | F>10 | 1 | 1 | 0.75 | 0.45 | 0.3 | 0.65 |
| | | F>30 | 1 | 1 | 0.2 | 0 | 0 | 0 |
| | | F>50 | 0.8 | 0.7 | 0.05 | 0 | 0 | 0 |
| | | F>100 | 0 | 0.05 | 0 | 0 | 0 | 0 |
| | 2000 | F>10 | 1 | 1 | 1 | 0.75 | 0.9 | 0.6 |
| | | F>30 | 1 | 1 | 0.6 | 0.15 | 0.1 | 0 |
| | | F>50 | 1 | 1 | 0.1 | 0 | 0.05 | 0 |
| | | F>100 | 1 | 1 | 0 | 0 | 0 | 0 |
| 0.5 | 500 | F>10 | 1 | 1 | 0.8 | 0.35 | 0.4 | 1.8 |
| | | F>30 | 1 | 1 | 0.35 | 0 | 0.05 | 0 |
| | | F>50 | 0.9 | 0.9 | 0 | 0 | 0 | 0 |
| | | F>100 | 0.2 | 0.3 | 0 | 0 | 0 | 0 |
| | 1000 | F>10 | 1 | 1 | 1 | 1 | 0.95 | 0.6 |
| | | F>30 | 1 | 1 | 0.8 | 0.5 | 0.15 | 0 |
| | | F>50 | 1 | 1 | 0.4 | 0.05 | 0 | 0 |
| | | F>100 | 1 | 1 | 0.05 | 0 | 0 | 0 |
| | 2000 | F>10 | 1 | 1 | 1 | 1 | 1 | 0.4 |
| | | F>30 | 1 | 1 | 1 | 0.9 | 0.9 | 0 |
| | | F>50 | 1 | 1 | 1 | 0.4 | 0.3 | 0 |
| | | F>100 | 1 | 1 | 0.35 | 0 | 0 | 0 |

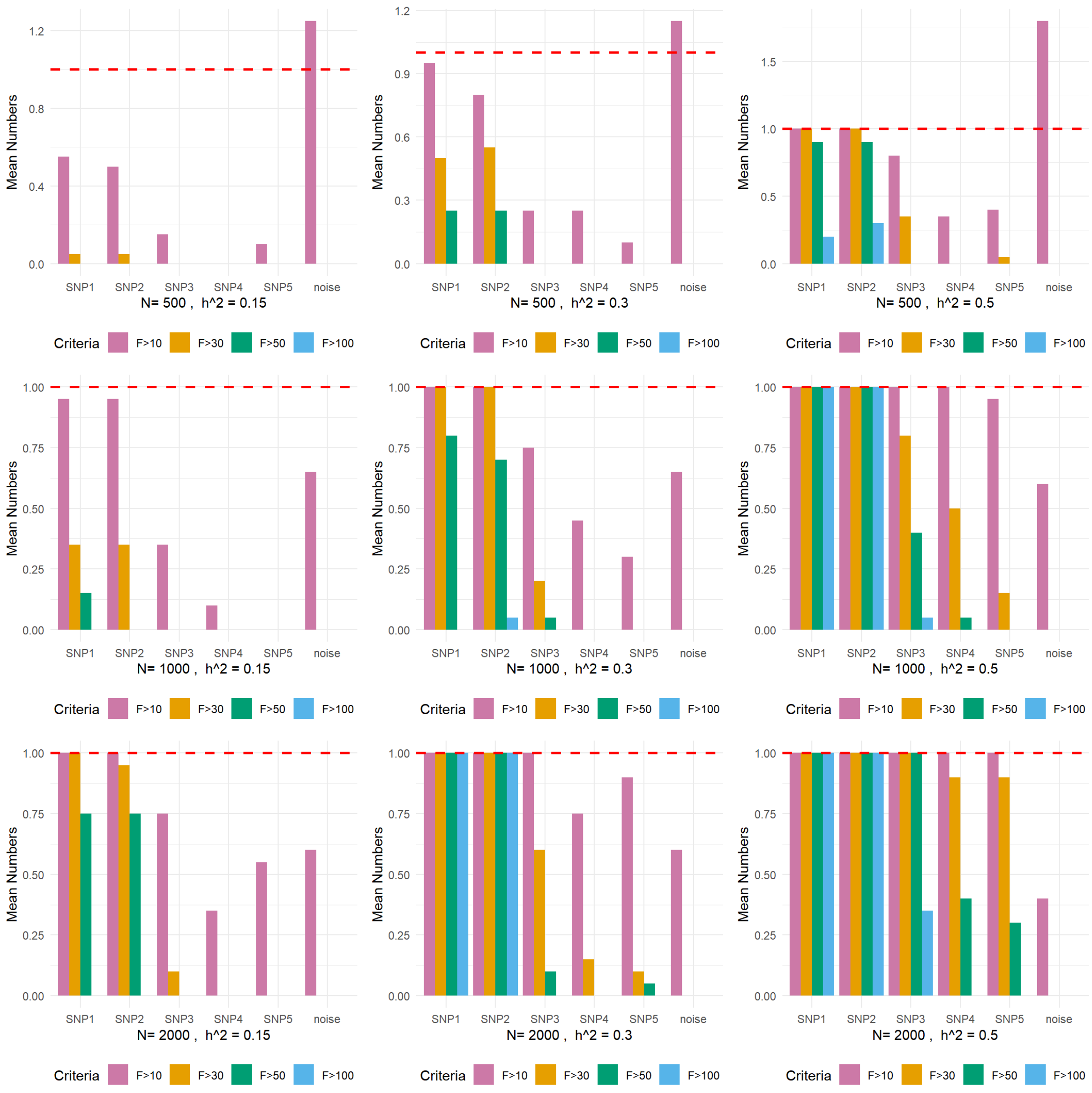


Figure S1: Mean numbers of being identified as major IV using different thresholds in 1,000 simulations.

#### 2 Boxplots of causal effect estimates for MR-SPLIT, 2SLS, and LIML out of 1000 simulation runs under different scenarios.

LIML and 2SLS both use half of the dataset to select IVs and the other half to get the estimation. In contrast, LIML_w and 2SLS_w use the whole dataset for IV selection and causal effect estimation. When using half data to select IVs and another half for causal estimation, both MR-SPLIT and LIML produce unbiased estimates, though the variance for LIML is larger than MR-SPLIT. However, when using the whole data for both IV selection and causal effect estimation, LIML_w and 2SLS_w generate biased causal effect estimation. In any either case, 2SLS yields biased effect estimates. This simulation demonstrates the issue of IV selection bias if it is not properly addressed.

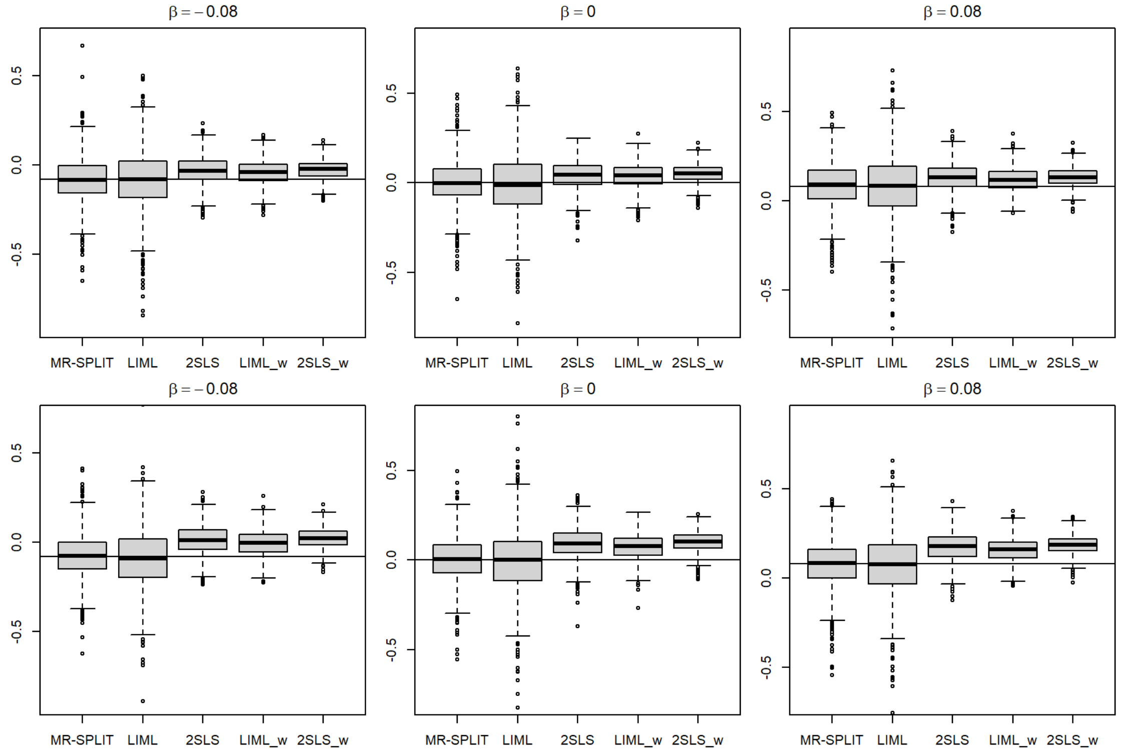


Figure S2: Boxplots of causal effect estimates ($\hat{\beta}$) under $h^2 = 0.15$ and confounding correlation $\rho = 0.1$ (top) and $\rho = 0.2$ (bottom).

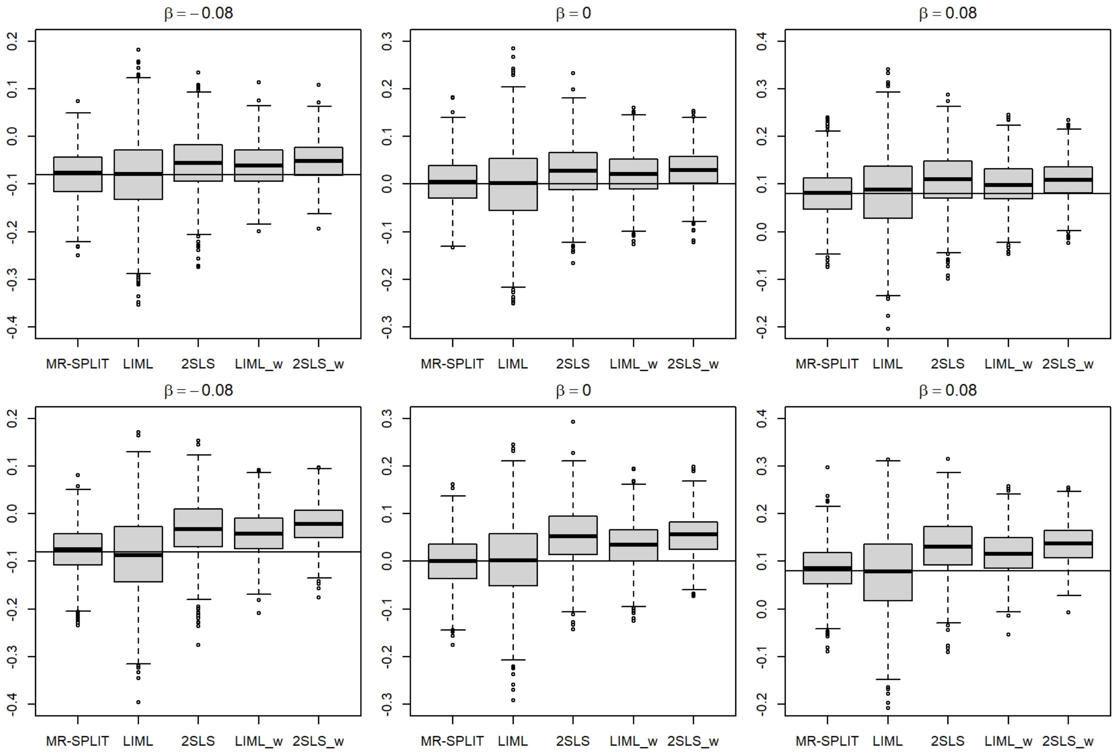


Figure S3: Boxplots of causal effect estimates ($\hat{\beta}$) under $h^2 = 0.3$ and confounding correlation $\rho = 0.1$ (top) and $\rho = 0.2$ (bottom).

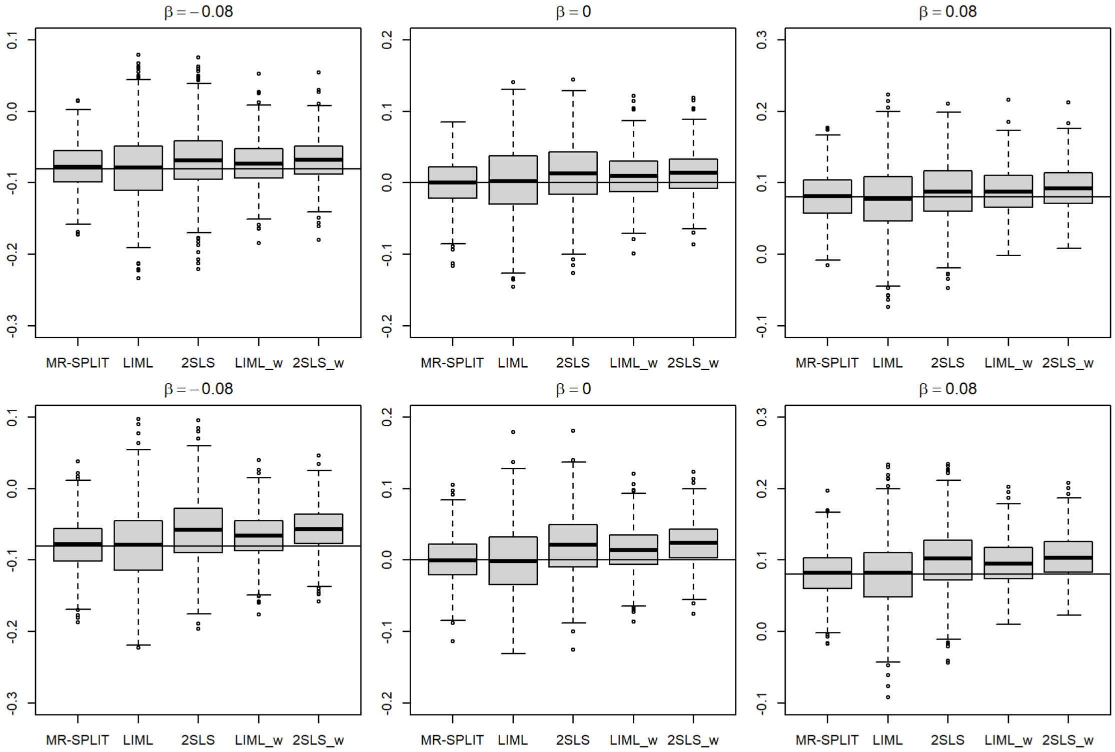


Figure S4: Boxplots of causal effect estimates ($\hat{\beta}$) under $h^2 = 0.5$ and confounding correlation $\rho = 0.1$ (top) and $\rho = 0.2$ (bottom).

#### 3 Boxplots of causal effect estimates for MR-SPLIT and CFMR out of 1000 simulation runs under different scenarios.

In nearly all scenarios, both CFMR and MR-SPLIT obtained estimates that are approximately unbiased. However, it is evident that MR-SPLIT consistently exhibits smaller variance compared to CFMR. This can also be seen in the comparison of RMSE (see Figure S11), where the RMSE of MR-SPLIT is always noticeably smaller than that of CFMR.

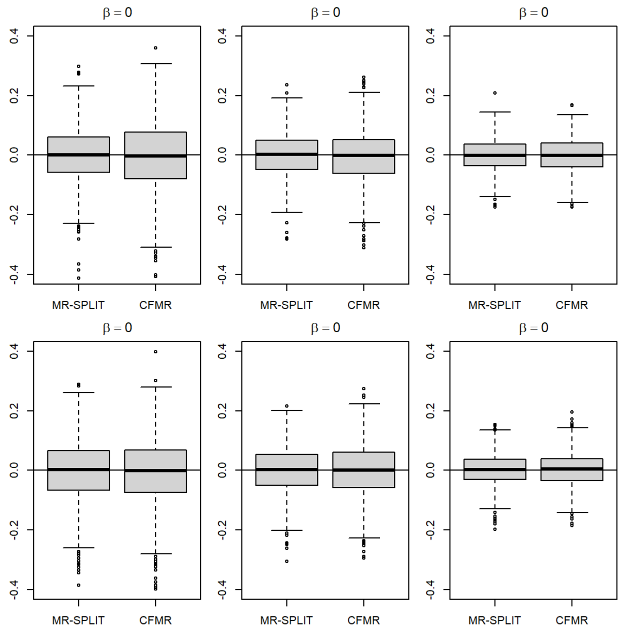


Figure S5: Boxplots of causal effect estimates ($\hat{\beta}$) when $h^2 = 0.15$ (left), $0.2$ (middle) , $0.3$ (right) and sample size $N = 1000$ in scenario I (top) and scenario II (bottom).

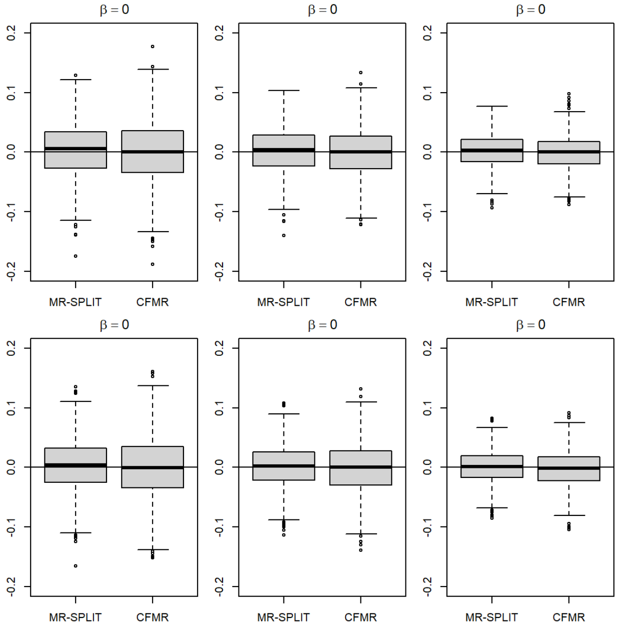


Figure S6: Boxplots of causal effect estimates ($\hat{\beta}$) when $h^2 = 0.15$ (left), 0.2 (middle) , 0.3 (right) and sample size $N = 3000$ in scenario I (top) and scenario II (bottom).

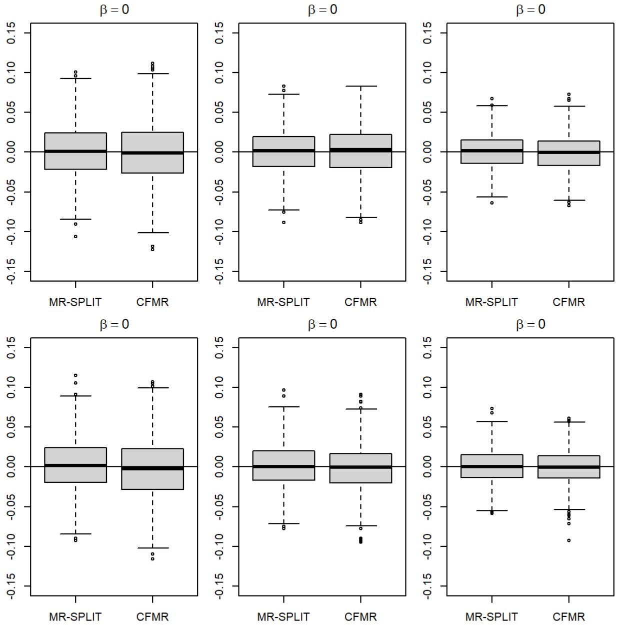


Figure S7: Boxplots of causal effect estimates ($\hat{\beta}$) when $h^2 = 0.15$ (left), 0.2 (middle) , 0.3 (right) and sample size $N = 5000$ in scenario I (top) and scenario II (bottom).

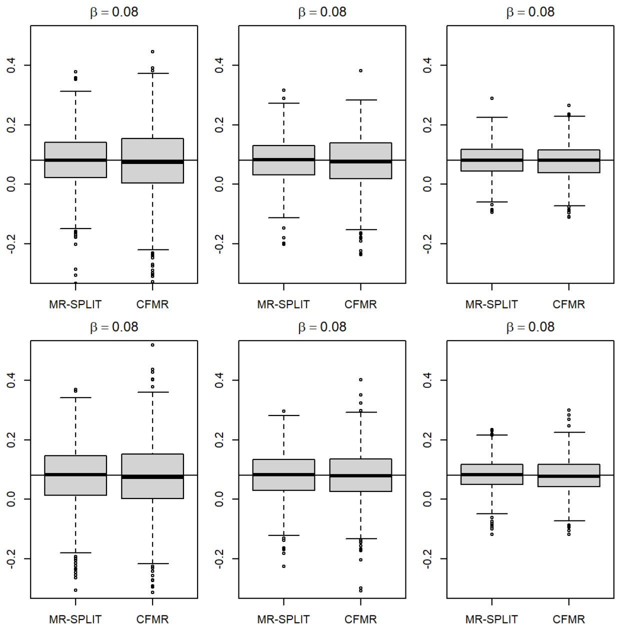


Figure S8: Boxplots of causal effect estimates ($\hat{\beta}$) when $h^2 = 0.15$ (left), $0.2$ (middle) , $0.3$ (right) and sample size $N = 1000$ in scenario I (top) and scenario II (bottom).

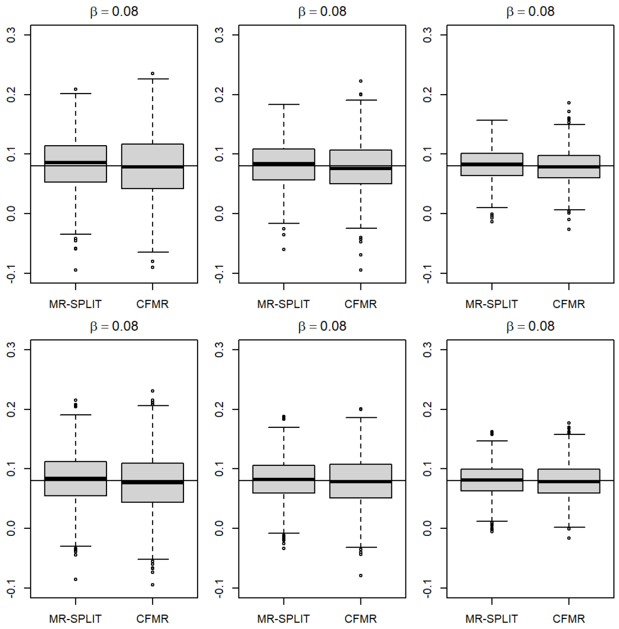


Figure S9: Boxplots of causal effect estimates ($\hat{\beta}$) when $h^2 = 0.15$ (left), $0.2$ (middle) , $0.3$ (right) and sample size $N = 3000$ in scenario I (top) and scenario II (bottom).

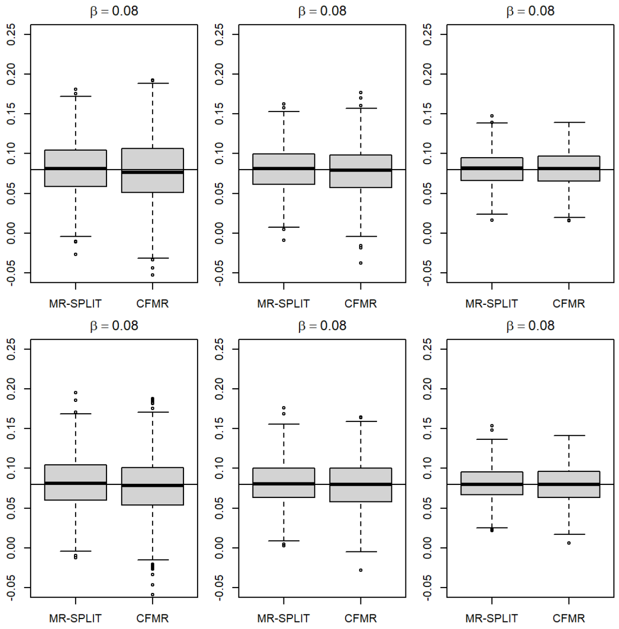


Figure S10: Boxplots of causal effect estimates ($\hat{\beta}$) when $h^2 = 0.15$ (left), 0.2 (middle) , 0.3 (right) and sample size $N = 5000$ in scenario I (top) and scenario II (bottom).

### 4 RMSE comparison between MR-SPLIT and CFMR out of 1000 simulation runs under different scenarios.

The RMSE of MR-SPLIT is always smaller than that of CFMR, especially under a small sample size (e.g., $N = 1000$), indicating the estimation efficiency and consistency of MR-SPLIT compared to CFMR.

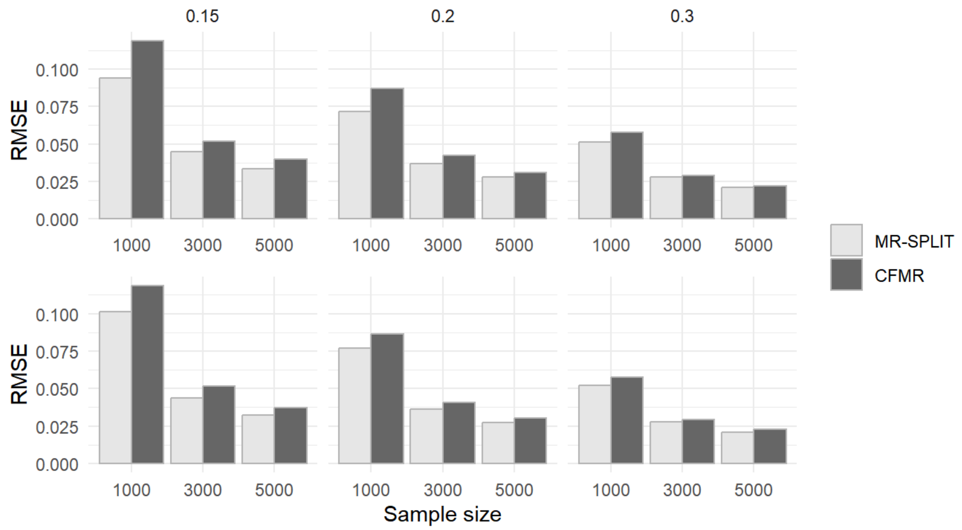


Figure S11: RNSE comparison between MR-SPLIT and CFMR in Scenario I (top) and II (bottom).

### 5 Additional type I error and power simulation results for the evaluation of multiple data splitting

Figure S12 shows the type I error out of 50 sample splits under $h^2 = 0.3$ and different sample sizes. When SNP heritability is significant and the sample size is relatively large (for instance, greater than 1000), the type I error stabilizes, even with a minimal number of sample splits.

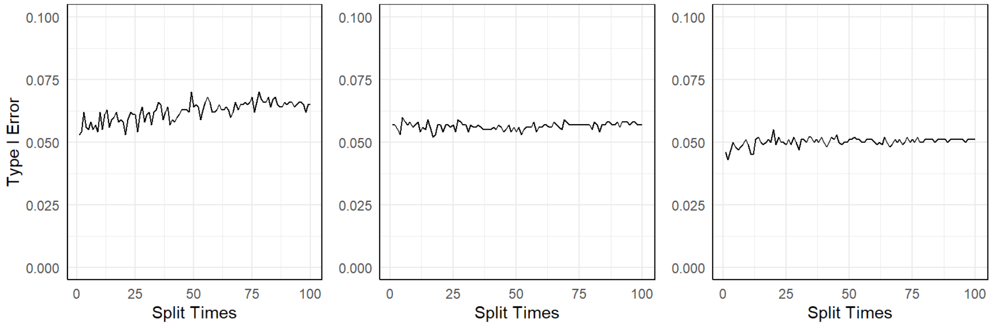


Figure S12: Type I error when $h^2 = 0.3$ and $N = 500$ (left), 1000 (middle), 2000 (right) out of 50 sample splits.

Figure S13 displays the empirical power under different sample sizes when $h^2 = 0.3$. Compared to scenarios where $h^2 = 0.15$ or 0.2, fewer splits are required to achieve optimal power. When the sample size is relatively small, for instance, $N = 500$, the power stabilizes after about 25 sample splits. As the sample size increases to 1000, a few sample splits are good enough to achieve stable power. The results suggest that in practice, one can lower the number of sample slits if the estimated SNP heritability for the exposure is strong and the sample size is large, to save computational time.

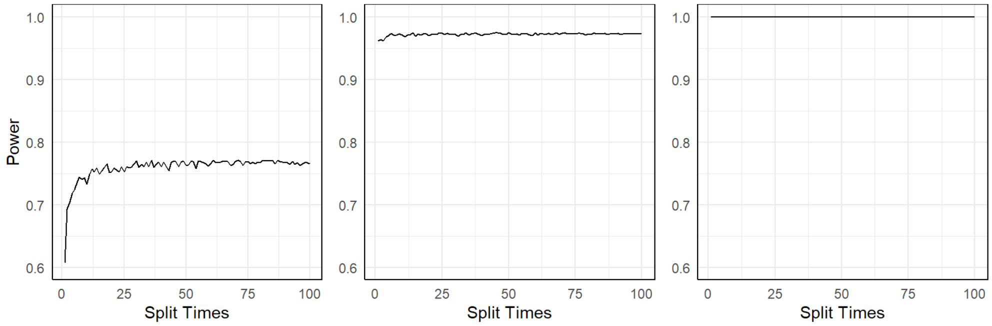


Figure S13: Power performance when $h^2 = 0.3$ and $N = 500$ (left), 1000 (middle), 2000 (right) out of 50 sample splits.

### 6 Additional results for the real data analysis

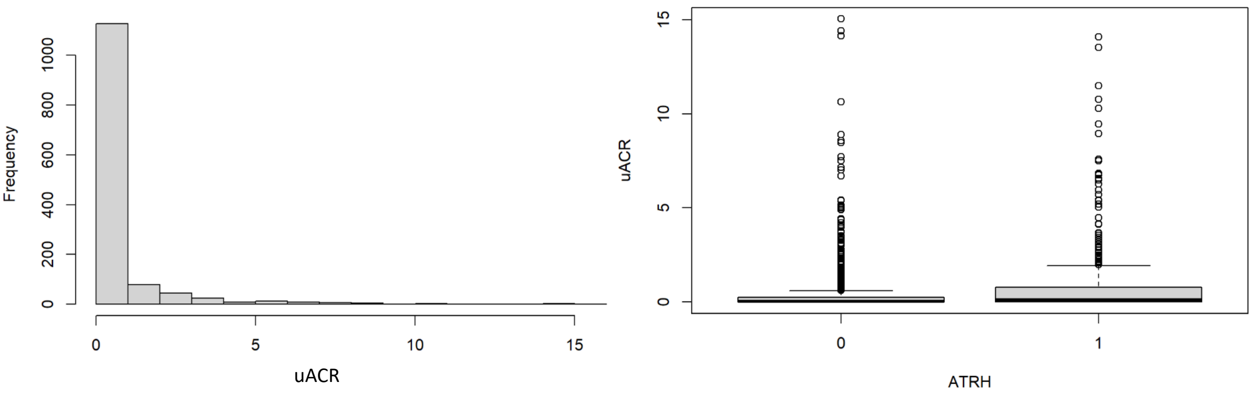


Figure S14: Distribution of uACR. Histogram (left figure) shows that the distribution of uACR is very skewed (to the right). It is difficult to see the difference of uACR in the two aTRH groups (right boxplots), indicating a transformation is needed for uACR.

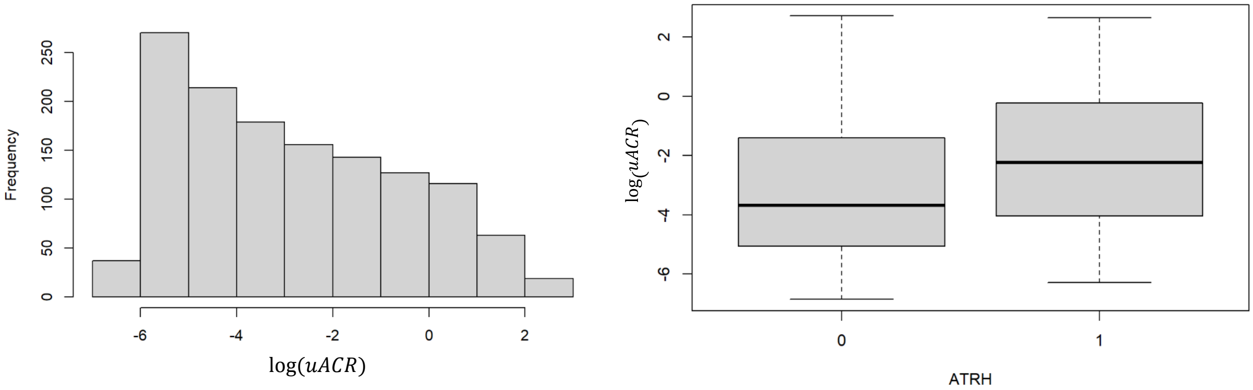


Figure S15: Distribution of log(uACR) (left figure) and the boxplots of log(uACR) in the two aTRH groups (right figure).

### 7 Proof of Theorem 1

For a given sample $\{X, Y, G\}$, the two stage IV model is defined as,

$$\begin{aligned} X &= G\alpha + \varepsilon_1 \\ Y &= X\beta + \varepsilon_2 \end{aligned} \tag{S.1}$$

where $(\varepsilon_1, \varepsilon_2)' \sim N(0, \sigma^2 \begin{pmatrix} 1 & \rho \\ \rho & 1 \end{pmatrix})$ and the correlation $\rho$ reflects the degree of confounding effect.

Suppose we split the data into two parts, $I_1 = \{X_1, Y_1, G_1\}$, and $I_2 = \{X_2, Y_2, G_2\}$. Each subset has equal sample size $N/2$, where $N$ is the total sample size. We first use sample $I_1$ to identify major and weak IVs, then use sample $I_2$ for causal inference. Suppose we have identified $p_1^{(1)}$ major IVs and $p_2^{(1)}$ weak IVs with the estimated effect size denoted as $\hat{\alpha}_1 = (\hat{\alpha}'_{1,M}, \hat{\alpha}'_{1,W})' \in \mathbb{R}^{p_1^{(1)} + p_2^{(1)}}$ when regressing exposure $X_1$ with the SNPs in $G_1$.

In sample $I_2$, MR-SPLIT combines the selected weak IVs into a new composite IV and uses it as an IV along with the major IVs:

$$\hat{G}_2 = (G_{2,M}, G_{2,W}\hat{\alpha}_{1,W}) \in \mathbb{R}^{\frac{N}{2} \times (p_1^{(1)}+1)} \tag{S.2}$$

Then, we can apply the stage one of 2SLS in sample $I_2$ using these IVs and get the estimates of the exposure in sample $I_2$:

$$\hat{X}_2 = \hat{G}_2(\hat{G}'_2\hat{G}_2)^{-1}\hat{G}'_2 X_2 = H_{\hat{G}_2} X_2,$$

where $H_X = X(X'X)^{-1}X'$ for any matrix $X$.

Similarly, we can also get the estimates of the exposure in sample $I_1$ by using sample $I_2$ to select the major and weak IVs:

$$\hat{X}_1 = \hat{G}_1(\hat{G}'_1\hat{G}_1)^{-1}\hat{G}'_1 X_1 = H_{\hat{G}_1} X_1$$

Let $\hat{X} = \begin{pmatrix} \hat{X}_1 \\ \hat{X}_2 \end{pmatrix}, Y = \begin{pmatrix} Y_1 \\ Y_2 \end{pmatrix}$. In stage two, we get the estimate of MR-SPLIT as

$$\begin{aligned} \hat{\beta} &= (\hat{X}'\hat{X})^{-1}\hat{X}'Y \\ &= \beta + (X'_1 H_{\hat{G}_1} X_1 + X'_2 H_{\hat{G}_2} X_2)^{-1}(X'_1 H_{\hat{G}_1}\varepsilon_{2,1} + X'_2 H_{\hat{G}_2}\varepsilon_{2,2}) \end{aligned}$$

and write $\varepsilon_2 = \begin{pmatrix} \varepsilon_{2,1} \\ \varepsilon_{2,2} \end{pmatrix}$.

For CFMR, it combines all selected IVs into a single IV. We use the subscript $C$ to denote variables used in CFMR:

$$\hat{G}_{2,C} = (G_{2,M}\hat{\alpha}_{1,M}, G_{2,W}\hat{\alpha}_{1,W}) = G_2\hat{\alpha}_1 \in \mathbb{R}^{n\times 1} \tag{S.3}$$

Similarly, in sample $I_1$, we combine $G_1$ and get:

$$\hat{G}_{1,C} = G_1\hat{\alpha}_2 \in \mathbb{R}^{n\times 1} \tag{S.4}$$

For CFMR, let $\hat{G}_C = \begin{pmatrix} \hat{G}_{1,C} \\ \hat{G}_{2,C} \end{pmatrix}, X = \begin{pmatrix} X_1 \\ X_2 \end{pmatrix}$. Apply 2SLS on $\{X, Y, \hat{G}_C\}$ we get

$$\begin{aligned} \hat{\beta}_C &= (X'H_{\hat{G}_C}X)^{-1}X'H_{\hat{G}_C}Y \\ &= \beta + (X'H_{\hat{G}_C}X)^{-1}X'H_{\hat{G}_C}\varepsilon_2 \end{aligned}$$

In the following, we will show that

$$var(\hat{\beta}) \le var(\hat{\beta}_C)$$

where $\hat{\beta}$ denotes the estimate by MR-SPLIT. Since $var(\hat{\beta}) = (X'_1 H_{\hat{G}_1} X_1 + X'_2 H_{\hat{G}_2} X_2)^{-1}\sigma^2$, $var(\hat{\beta}_C) = (X'H_{\hat{G}_C}X)^{-1}\sigma^2$, to prove $var(\hat{\beta}) \le var(\hat{\beta}_C)$, we need to show

$$\begin{aligned} & X'_1 H_{\hat{G}_1} X_1 + X'_2 H_{\hat{G}_2} X_2 \ge X'H_{\hat{G}_C}X \\ \iff\ & X' \begin{pmatrix} H_{\hat{G}_1} & \\ & H_{\hat{G}_2} \end{pmatrix} X \ge X'H_{\hat{G}_C}X \\ \iff\ & X' \left( \begin{pmatrix} H_{\hat{G}_1} & \\ & H_{\hat{G}_2} \end{pmatrix} - H_{\hat{G}_C} \right) X \ge 0 \end{aligned}$$

Hence, it is sufficient to show

$$\begin{pmatrix} H_{\hat{G}_1} & \\ & H_{\hat{G}_2} \end{pmatrix} - H_{\hat{G}_C} \succeq 0 \tag{S.5}$$

where for any matrix $X$, $X \succeq 0$ means it is positive semi-definite.

Recall that $\hat{G}_C = \begin{pmatrix} \hat{G}_{1,C} \\ \hat{G}_{2,C} \end{pmatrix}$,

$$H_{\hat{G}_C} = \hat{G}_C(\hat{G}_C'\hat{G}_C)^{-1}\hat{G}_C' = \frac{1}{\hat{G}_{1,C}'\hat{G}_{1,C} + \hat{G}_{2,C}'\hat{G}_{2,C}} \begin{pmatrix} \hat{G}_{1,C}\hat{G}_{1,C}' & \hat{G}_{1,C}\hat{G}_{2,C}' \\ \hat{G}_{2,C}\hat{G}_{1,C}' & \hat{G}_{2,C}\hat{G}_{2,C}' \end{pmatrix}$$

Let $a = \hat{G}_{1,C}'\hat{G}_{1,C} + \hat{G}_{2,C}'\hat{G}_{2,C} \in \mathbb{R}$, it remains to show

$$\begin{pmatrix} H_{\hat{G}_1} - \frac{\hat{G}_{1,C}\hat{G}_{1,C}'}{a} & -\frac{\hat{G}_{1,C}\hat{G}_{2,C}'}{a} \\ -\frac{\hat{G}_{2,C}\hat{G}_{1,C}'}{a} & H_{\hat{G}_2} - \frac{\hat{G}_{2,C}\hat{G}_{2,C}'}{a} \end{pmatrix} \succeq 0 \tag{S.6}$$

From Eq. S.2 Eq. S.3 and Eq. S.4, we can get

$$\begin{aligned}
H_{\hat{G}_2} =& \begin{pmatrix} G_{2,M} & G_{2,W}\hat{\alpha}_{1,W} \end{pmatrix} \left(\hat{G}_2'\hat{G}_2\right)^{-1} \begin{pmatrix} G_{2,M}' \\ \hat{\alpha}_{1,W}'G_{2,W}' \end{pmatrix} \\
=& \begin{pmatrix} G_{2,M} & G_{2,W}\hat{\alpha}_{1,W} \end{pmatrix} \begin{pmatrix} A_2 & B_2 \\ C_2 & D_2 \end{pmatrix} \begin{pmatrix} G_{2,M}' \\ \hat{\alpha}_{1,W}'G_{2,W}' \end{pmatrix} \\
=& G_{2,M}A_2G_{2,M}' + G_{2,W}\hat{\alpha}_{1,W}C_2G_{2,M}' + G_{2,M}B_2\hat{\alpha}_{1,W}'G_{2,W}' + G_{2,W}\hat{\alpha}_{1,W}D_2\hat{\alpha}_{1,W}'G_{2,W}' \\
\frac{\hat{G}_{2,C}\hat{G}_{2,C}'}{a} =& \frac{1}{a} \begin{pmatrix} G_{2,M} & G_{2,W} \end{pmatrix} \begin{pmatrix} \hat{\alpha}_{1,M}\hat{\alpha}_{1,M}' & \hat{\alpha}_{1,M}\hat{\alpha}_{1,W}' \\ \hat{\alpha}_{1,W}\hat{\alpha}_{1,M}' & \hat{\alpha}_{1,W}\hat{\alpha}_{1,W}' \end{pmatrix} \begin{pmatrix} G_{2,M}' \\ G_{2,W}' \end{pmatrix} \\
=& \frac{1}{a}(G_{2,M}\hat{\alpha}_{1,M}\hat{\alpha}_{1,M}'G_{2,M}' + G_{2,W}\hat{\alpha}_{1,W}\hat{\alpha}_{1,M}'G_{2,M}' \\
&+ G_{2,M}\hat{\alpha}_{1,M}\hat{\alpha}_{1,W}'G_{2,W}' + G_{2,W}\hat{\alpha}_{1,W}\hat{\alpha}_{1,W}'G_{2,W}')
\end{aligned} \tag{S.7}$$

Therefore,

$$\begin{aligned}
H_{\hat{G}_2} - \frac{\hat{G}_{2,C}\hat{G}_{2,C}'}{a} =& G_{2,M}\left(A_2 - \frac{\hat{\alpha}_{1,M}\hat{\alpha}_{1,M}'}{a}\right)G_{2,M}' + G_{2,W}\hat{\alpha}_{1,W}\left(C_2 - \frac{\hat{\alpha}_{1,M}'}{a}\right)G_{2,M}' \\
&+ G_{2,M}\left(B_2 - \frac{\hat{\alpha}_{1,M}}{a}\right)\hat{\alpha}_{1,W}'G_{2,W}' + G_{2,W}\hat{\alpha}_{1,W}(D_2 - \frac{1}{a})\hat{\alpha}_{1,W}'G_{2,W}' \\
=& \begin{pmatrix} G_{2,M} & G_{2,W}\hat{\alpha}_{1,W} \end{pmatrix} \begin{pmatrix} A_2 - \frac{\hat{\alpha}_{1,M}\hat{\alpha}_{1,M}'}{a} & B_2 - \frac{\hat{\alpha}_{1,M}}{a} \\ C_2 - \frac{\hat{\alpha}_{1,M}'}{a} & D_2 - \frac{1}{a} \end{pmatrix} \begin{pmatrix} G_{2,M}' \\ \hat{\alpha}_{1,W}'G_{2,W}' \end{pmatrix} \\
=& \hat{G}_2Q_4\hat{G}_2'
\end{aligned} \tag{S.8}$$

Similarly,

$$\begin{aligned}
H_{\hat{G}_1} - \frac{\hat{G}_{1,C}\hat{G}_{1,C}'}{a} =& \begin{pmatrix} G_{1,M} & G_{1,W}\hat{\alpha}_{2,W} \end{pmatrix} \begin{pmatrix} A_1 - \frac{\hat{\alpha}_{2,M}\hat{\alpha}_{2,M}'}{a} & B_1 - \frac{\hat{\alpha}_{2,M}}{a} \\ C_1 - \frac{\hat{\alpha}_{2,M}'}{a} & D_1 - \frac{1}{a} \end{pmatrix} \begin{pmatrix} G_{1,M}' \\ \hat{\alpha}_{2,W}'G_{1,W}' \end{pmatrix} \\
=& \hat{G}_1Q_1\hat{G}_1'
\end{aligned} \tag{S.9}$$

Easily, we can also get

$$\begin{aligned}
-\frac{\hat{G}_{1,C}\hat{G}_{2,C}'}{a} =& -\frac{1}{a} \begin{pmatrix} G_{1,M} & G_{1,W}\hat{\alpha}_{2,W} \end{pmatrix} \begin{pmatrix} \hat{\alpha}_{2,M}\hat{\alpha}_{1,M} & \hat{\alpha}_{2,M} \\ \hat{\alpha}_{1,M}' & 1 \end{pmatrix} \begin{pmatrix} G_{2,M}' \\ \hat{\alpha}_{1,W}'G_{2,W}' \end{pmatrix} \\
=& -\frac{1}{a}\hat{G}_1Q_2\hat{G}_2'
\end{aligned} \tag{S.10}$$

$$\begin{aligned}
-\frac{\hat{G}_{2,C}\hat{G}_{1,C}'}{a} =& -\frac{1}{a} \begin{pmatrix} G_{2,M} & G_{2,W}\hat{\alpha}_{1,W} \end{pmatrix} \begin{pmatrix} \hat{\alpha}_{1,M}\hat{\alpha}_{2,M} & \hat{\alpha}_{1,M} \\ \hat{\alpha}_{2,M}' & 1 \end{pmatrix} \begin{pmatrix} G_{1,M}' \\ \hat{\alpha}_{2,W}'G_{1,W}' \end{pmatrix} \\
=& -\frac{1}{a}\hat{G}_2Q_3\hat{G}_1'
\end{aligned} \tag{S.11}$$

Apply Eq.S.8, Eq.S.9, Eq.S.10, Eq.S.11 to Eq. S.6, we have

$$\begin{pmatrix} \hat{G}_1 & \hat{G}_2 \end{pmatrix} \begin{pmatrix} Q_1 & Q_2 \\ Q_3 & Q_4 \end{pmatrix} \begin{pmatrix} \hat{G}_1' \\ \hat{G}_2' \end{pmatrix} \succeq 0 \tag{S.12}$$

We now only need to show that

$$\begin{pmatrix} Q_1 & Q_2 \\ Q_3 & Q_4 \end{pmatrix} \succeq 0. \tag{S.13}$$

We first show that

$$Q_4 = \begin{pmatrix} A_2 - \frac{\hat{\alpha}_{1,M}\hat{\alpha}_{1,M}'}{a} & B_2 - \frac{\hat{\alpha}_{1,M}}{a} \\ C_2 - \frac{\hat{\alpha}_{1,M}'}{a} & D_2 - \frac{1}{a} \end{pmatrix} \succ 0. \tag{S.14}$$

Recall that

$$\begin{aligned} \begin{pmatrix} A_2 & B_2 \\ C_2 & D_2 \end{pmatrix} &= (\hat{G}_2'\hat{G}_2)^{-1} \\ &= \begin{pmatrix} G_{2,M}'G_{2,M} & G_{2,M}'G_{2,W}\hat{\alpha}_{1,W} \\ \hat{\alpha}_{1,W}'G_{2,W}'G_{2,M} & \hat{\alpha}_{1,W}'G_{2,W}'G_{2,W}\hat{\alpha}_{1,W} \end{pmatrix}^{-1} \end{aligned} \tag{S.15}$$

Thus,

$$\begin{aligned} A_2 &= (G_{2,M}'G_{2,M})^{-1} + \frac{(G_{2,M}'G_{2,M})^{-1}(G_{2,M}'G_{2,W}\hat{\alpha}_{1,W}\hat{\alpha}_{1,W}'G_{2,W}'G_{2,M})(G_{2,M}'G_{2,M})^{-1}}{\hat{\alpha}_{1,W}'G_{2,W}'(I - H_{G_{2,M}})G_{2,W}\hat{\alpha}_{1,W}}, \\ B_2 &= -\frac{(G_{2,M}'G_{2,M})^{-1}G_{2,M}'G_{2,W}\hat{\alpha}_{1,W}}{\hat{\alpha}_{1,W}'G_{2,W}'(I - H_{G_{2,M}})G_{2,W}\hat{\alpha}_{1,W}}, \\ C_2 &= -\frac{\hat{\alpha}_{1,W}'G_{2,W}'G_{2,W}(G_{2,M}'G_{2,M})^{-1}}{\hat{\alpha}_{1,W}'G_{2,W}'(I - H_{G_{2,M}})G_{2,W}\hat{\alpha}_{1,W}}, \\ D_2 &= \frac{1}{\hat{\alpha}_{1,W}'G_{2,W}'(I - H_{G_{2,M}})G_{2,W}\hat{\alpha}_{1,W}}. \end{aligned}$$

Then,

$$\begin{aligned} D_2 - \frac{1}{a} =& \frac{1}{\hat{\alpha}_{1,W}'G_{2,W}'(I - H_{G_{2,M}})G_{2,W}\hat{\alpha}_{1,W}} - \frac{1}{\hat{G}_{1,C}'\hat{G}_{1,C} + \hat{G}_{2,C}'\hat{G}_{2,C}} \\ =& \frac{1}{\hat{\alpha}_{1,W}'G_{2,W}'(I - H_{G_{2,M}})G_{2,W}\hat{\alpha}_{1,W}} \\ &- \frac{1}{\hat{G}_{1,C}'\hat{G}_{1,C} + (G_{2,M}\hat{\alpha}_{1,M} + G_{2,W}\hat{\alpha}_{1,W})'(G_{2,M}\hat{\alpha}_{1,M} + G_{2,W}\hat{\alpha}_{1,W})} \end{aligned}$$

Since

$$\begin{aligned} &(G_{2,M}\hat{\alpha}_{1,M} + G_{2,W}\hat{\alpha}_{1,W})'(G_{2,M}\hat{\alpha}_{1,M} + G_{2,W}\hat{\alpha}_{1,W}) - \hat{\alpha}_{1,W}'G_{2,W}'(I - H_{G_{2,M}})G_{2,W}\hat{\alpha}_{1,W} \\ =&(G_{2,M}\hat{\alpha}_{1,M} + H_{G_{2,M}}G_{2,W}\hat{\alpha}_{1,W})'(G_{2,M}\hat{\alpha}_{1,M} + H_{G_{2,M}}G_{2,W}\hat{\alpha}_{1,W}) > 0, \end{aligned} \tag{S.16}$$

we get $D_2 - \frac{1}{a} > 0$.
To prove (S.14), it is sufficient to show[? ]

$$\begin{aligned} &A_2 - \frac{\hat{\alpha}_{1,M}\hat{\alpha}_{1,M}'}{a} - (B_2 - \frac{\hat{\alpha}_{1,M}}{a})(D_2 - \frac{1}{Q})^{-1}(C_2 - \frac{\hat{\alpha}_{1,M}'}{a}) \succ 0 \\ \Longleftrightarrow &(A_2 - \frac{\hat{\alpha}_{1,M}\hat{\alpha}_{1,M}'}{a})(D_2 - \frac{1}{a}) - (B_2 - \frac{\hat{\alpha}_{1,M}}{a})(C_2 - \frac{\hat{\alpha}_{1,M}'}{a}) \succ 0 \\ \Longleftrightarrow &(a - \frac{1}{D_2})\tilde{A}_2^{-1} - \hat{\alpha}_{1,M}\hat{\alpha}_{1,M}' + \tilde{A}_2^{-1}\tilde{B}_2\tilde{C}_2\tilde{A}_2^{-1} - \hat{\alpha}_{1,M}\tilde{C}_2\tilde{A}_2^{-1} - \tilde{A}_2^{-1}\tilde{B}_2\hat{\alpha}_{1,M}' \succ 0 \end{aligned} \tag{S.17}$$

where $\begin{pmatrix} \tilde{A}_2 & \tilde{B}_2 \\ \tilde{C}_2 & \tilde{D}_2 \end{pmatrix} = \hat{G}_2'\hat{G}_2$. The left side of Eq. S.17 can be obtained as $(a - \frac{1}{D_2})\tilde{A}_2^{-1} + (\tilde{A}_2^{-1}B_2 - \hat{\alpha}_{1,M})(\tilde{A}_2^{-1}B_2 - \hat{\alpha}_{1,M})'$, which is easy to verify to be a positive definite matrix.
To prove Eq. S.13, now we only need to prove

$$Q_1 - Q_2Q_4^{-1}Q3 \succeq 0. \tag{S.18}$$

From Eq. S.14, we have

$$Q_4^{-1} = \left( (\hat{G}_2'\hat{G}_2)^{-1} - \frac{1}{a} \begin{pmatrix} \hat{\alpha}_{1,M} \\ 1 \end{pmatrix} \begin{pmatrix} \hat{\alpha}_{1,M}' & 1 \end{pmatrix} \right)^{-1}$$
$$= \left( (\hat{G}_2'\hat{G}_2)^{-1} - \frac{1}{a} b_2 b_2' \right)^{-1}$$
$$= \hat{G}_2'\hat{G}_2 - \frac{\hat{G}_2'\hat{G}_2 b_2 b_2' \hat{G}_2'\hat{G}_2}{b_2'\hat{G}_2'\hat{G}_2 b_2 - a}, \tag{S.19}$$

where $b_2 = \left(\hat{\alpha}_{1,M}', 1\right)'$, and the third equation utilizes the Woodbury matrix identity. Similarly, let $b_1 = \left(\hat{\alpha}_{2,M}', 1\right)'$,

$$Q_1 = (\hat{G}_1'\hat{G}_1)^{-1} - \frac{1}{a} b_1 b_1', \tag{S.20}$$
$$Q_2 = -\frac{1}{a} b_1 b_2', \tag{S.21}$$
$$Q_3 = -\frac{1}{a} b_2 b_1'. \tag{S.22}$$

Substituting Eq. S.19, Eq. S.20, Eq. S.21, and Eq. S.22 into Eq. S.18, we get

$$(\hat{G}_1'\hat{G}_1)^{-1} - \frac{1}{a} b_1 b_1' - \frac{1}{a^2} b_1 b_2' (\hat{G}_2'\hat{G}_2 - \frac{\hat{G}_2'\hat{G}_2 b_2 b_2' \hat{G}_2'\hat{G}_2}{b_2'\hat{G}_2'\hat{G}_2 b_2 - a}) b_2 b_1' \succeq 0$$
$$\Longleftrightarrow (\hat{G}_1'\hat{G}_1)^{-1} - b_1 (\frac{1}{a} + \frac{1}{a^2} b_2' (\hat{G}_2'\hat{G}_2 - \frac{\hat{G}_2'\hat{G}_2 b_2 b_2' \hat{G}_2'\hat{G}_2}{b_2'\hat{G}_2'\hat{G}_2 b_2 - a}) b_2) b_1' \succeq 0$$
$$\Longleftrightarrow (\hat{G}_1'\hat{G}_1)^{-1} - b_1 (\frac{1}{a} + \frac{\hat{G}_{2,C}'\hat{G}_{2,C}}{a^2} - \frac{(\hat{G}_{2,C}'\hat{G}_{2,C})^2}{a^2 \hat{G}_{2,C}'\hat{G}_{2,C} - a^3}) b_1' \succeq 0$$
$$\Longleftrightarrow (\hat{G}_1'\hat{G}_1)^{-1} - \frac{1}{\hat{G}_{1,C}'\hat{G}_{1,C}} b_1 b_1' \succeq 0 \tag{S.23}$$

Eq. S.23 has a very similar structure as Eq. S.14. It can be written as

$$\begin{pmatrix} A_1 - \frac{\hat{\alpha}_{2,M}\hat{\alpha}_{2,M}'}{\hat{G}_{1,C}'\hat{G}_{1,C}} & B_1 - \frac{\hat{\alpha}_{2,M}}{\hat{G}_{1,C}'\hat{G}_{1,C}} \\ C_1 - \frac{\hat{\alpha}_{2,M}'}{\hat{G}_{1,C}'\hat{G}_{1,C}} & D_1 - \frac{1}{\hat{G}_{1,C}'\hat{G}_{1,C}} \end{pmatrix} \succeq 0, \tag{S.24}$$

which can be easily verified. This completes the proof of the theorem.
